# Constitutive expression of a fluorescent protein reports the size of live human cells

**DOI:** 10.1101/587162

**Authors:** Daniel F. Berenson, Evgeny Zatulovskiy, Shicong Xie, Jan M. Skotheim

## Abstract

Cell size is intimately related to cell physiology because it sets the geometric scale of organelles and biosynthesis. A number of methods exist to measure different aspects of size of individual cells, but each has significant drawbacks. Here, we present an alternative method to measure the size of single human cells using a nuclear localized fluorescent protein expressed from a constitutive promoter. We validate this method by comparing it to several established cell size measurement strategies, including flow cytometry optical scatter, total protein dyes, and quantitative phase microscopy. We directly compare our fluorescent protein measurement to the commonly used measurement of nuclear volume and show that our measurements are more robust and less dependent on image segmentation. We apply our method to examine how cell size impacts the cell division cycle, which reaffirms the importance of G1/S size control. Importantly, combining our size reporter with fluorescent labeling of a different protein in a different color channel allows measurement of concentration dynamics using simple widefield fluorescence imaging. Thus, we expect our method will be of use to other researchers interested in the topics of cell size control and, more broadly, how dynamically changing protein concentrations control cell fates.

## INTRODUCTION

Cell size has an important effect on cellular physiology through its influence on biosynthesis, mitochondrial efficiency, and hormone secretion (Figure 1A) (Smith, 1971; Pende *et al.*, 2000; Miettinen and Björklund, 2016). Variation in cell size is one of the most noticeable differences between cells of different type and function (Ginzberg *et al.*, 2015). However, this size variation is rarely emphasized by researchers, possibly because it has been difficult to measure. While a growing number of methods provide size information, either as the experiment’s primary objective or as an additional parameter during experiments designed for other purposes, there is a need for more convenient methods for measuring size during live-cell microscopy (Table 1).

**Table 1.** Comparison of methods to measure cell size.

| Method | Measures | Does not require custom equipment | High throughput | Compatible with live cell imaging | References |
| --- | --- | --- | --- | --- | --- |
| Flow cytometry forward or side scatter (FSC or SSC) | FSC correlates with cell volume<br>SSC correlates with cell volume and granularity | ✓ | ✓ ✓ | ✗ | Shapiro, *Practical Flow Cytometry* |
| Coulter counter | Cell volume | ✓ | ✓ ✓ | ✗ | Tzur *et al.*, 2009 |
| Estimation of nuclear volume by microscopy | Nuclear area, then estimate nuclear volume, correlated with cell size | ✓ | ✓ | ✓ | Liu *et al.*, 2018; Ginzberg *et al.*, 2018 |
| Succinimidyl ester fluorescent protein dye (e.g., CFSE) | Cellular total protein content | ✓ | ✓ | ✗ | Kafri *et al.*, 2013 |
| Confocal stack | Cell volume | ✓ | ✗ | ✓ | Bajcsy *et al.*, 2015 |
| Quantitative phase microscopy | Dry mass | ✓ | ✗ | ✓ | Barer, 1952; Kafri *et al.*, 2013; Zangle and Teitell, 2015 |
| Microchannel resonator | Buoyant mass | ✗ | ✗ | ✓ | Godin *et al.*, 2010; Son *et al.*, 2012; Son *et al.*, 2015 |
| Fluorescence exclusion | Cell volume | ✗ | ✓ | ✓ | Bottier *et al.*, 2011; Zlotek-Zlotkiewicz *et al.*, 2015; Cadart *et al.*, 2018 |
| Channel assisted reshaping | Cell volume | ✗ | ✓ | ✓ | Varsano *et al.*, 2017 |
| Constitutively expressed fluorescent protein | Promoter expression correlated with cell size | ✓ | ✓ | ✓ | DiTalia *et al.*, 2007; this paper. |

**Figure 1.**
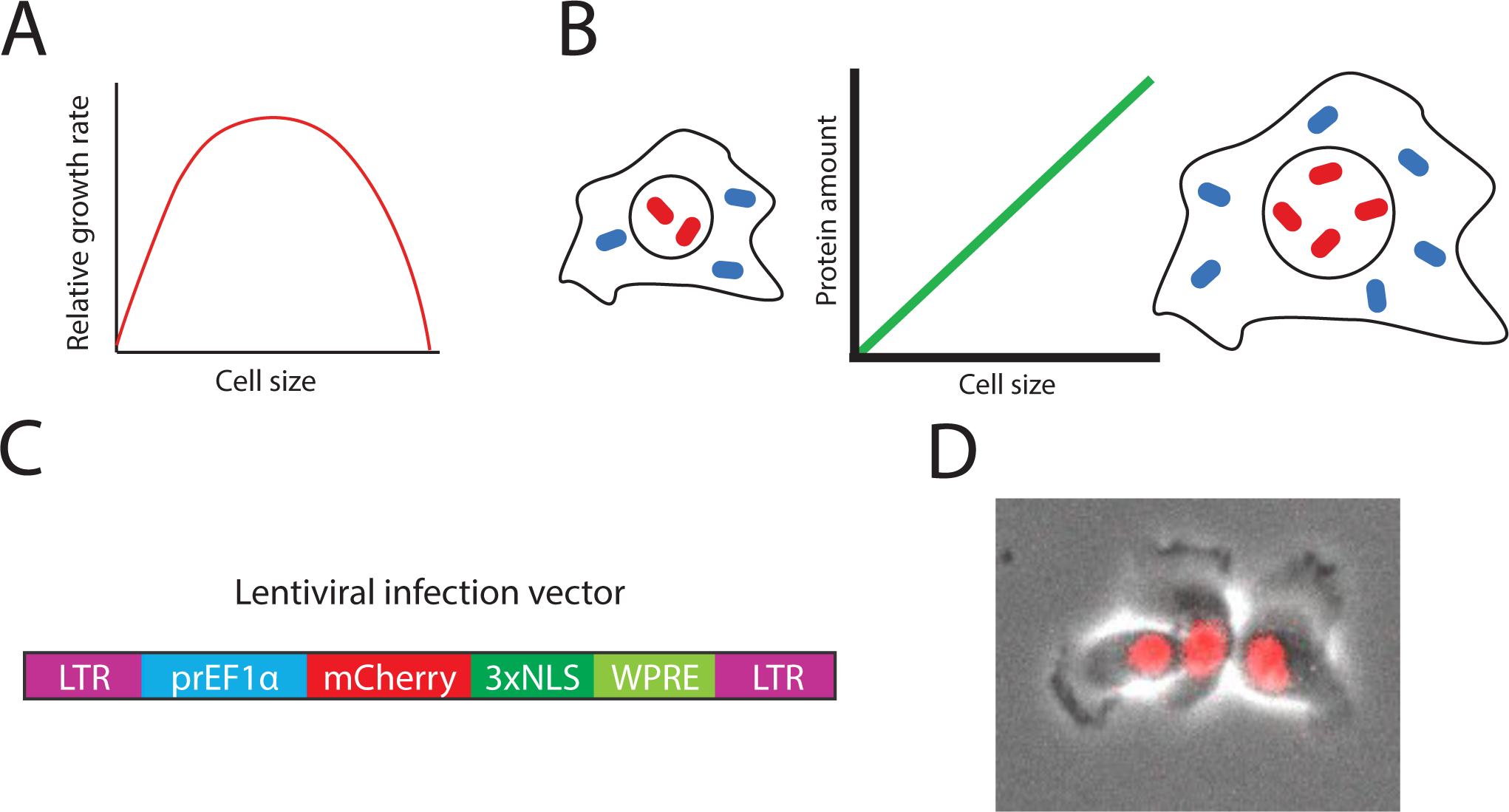
Measuring cell size using a constitutively expressed fluorescent protein. (A) Single cell growth rate has an optimum as a function of cell size (Miettinen and Björklund, 2016). (B) Principle of size reporter: the amount of constitutively expressed proteins increases in proportion to cell size. (C) Schematic of the lentiviral infection vector. LTR denotes long terminal repeats, *prEF1α* denotes 1kb of the *EF1α* promoter, NLS denotes nuclear localization sequence, and WPRE denotes a woodchuck posttranscriptional regulatory element that boosts expression (Zufferey *et al.*, 1999). (D) Representative composite phase and fluorescence image of HMEC cells expressing mCherry-NLS from an *EF1α* promoter.

Further complicating accurate measurement of cell size is the ambiguity as to what exactly “size” means. In general, researchers mean one of three things: volume, dry mass, or protein content. Different techniques exist to measure each of these parameters, but all three mostly correlate and are thought to reflect size. That is, cells of a given type in a particular condition have constant ratios of mass to volume and of protein content to mass. However, some cells - such as mitotic cells, chondrocytes, and cell-cycle-blockaded budding yeast - dilute their dry mass with water so it is important to understand which parameter a particular technique is measuring (Cooper *et al.*, 2013; Son *et al.*, 2015; Zlotek-Zlotkiewicz *et al.*, 2015; Neurohr *et al.*, 2019).

The gold standard for cell volume measurements is considered to be the Coulter counter, which flows cells through a measurement chamber where they displace an isotonic solution to cause a volume-proportional change in electrical impedance (Gregg and Steidley, 1965). Coulter counters measure cell number and size in high throughput, but do not provide any additional information (such as amount of a protein of interest) and cannot make multiple measurements on the same cell over time (Tzur *et al.*, 2009). A related single time point high-throughput technique is flow cytometry, in which cells pass one by one through a series of lasers. In combination with immunofluorescence techniques labeling individual protein species, flow cytometry can be used to collect multi-dimensional, high-throughput data. Flow cytometers also measure two optical properties, forward scatter (FSC) and side scatter (SSC), that reflect cell size, shape, and granularity. Although commonly used as a proxy for size, evidence for a quantitative, linear relationship between FSC and cell size is sparse, and the relationship depends on cell type and flow cytometer machine and settings (Holme *et al.*, 1988; Tzur *et al.*, 2011; Shapiro, 2018). Calibration of size estimates from cytometry scatter is often done by comparison to fluorescent dyes that stain total cellular protein. These dyes, CFSE and SE-Alexa647, use reactive succinimidyl ester groups to react with free amino groups on proteins, so that the amount of staining corresponds to total cellular protein (Kafri *et al.*, 2013; Shapiro, 2018). Importantly, these dyes do not provide quantitative information on total protein when used in live-cell imaging.

Live-cell microscopy provides a unique and rich source of single-cell data on cellular processes through time. Live-cell studies have been particularly impactful for studying the cell cycle, where molecular signaling and the passage of time both contribute to progression through cell cycle phases (Doncic *et al.*, 2011; Doncic and Skotheim, 2013; Spencer *et al.*, 2013; Atay *et al.*, 2016; Cappell *et al.*, 2016; Barr *et al.*, 2017; Schwarz *et al.*, 2018). Since cell size changes over time in growing cells and has been shown to be an important driver of cell cycle transitions, it is particularly important to accurately measure cell size in cell cycle experiments (DiTalia *et al.*, 2007; Schmoller *et al.*, 2015). Unfortunately, such studies, particularly in the irregularly shaped animal cells, have been limited by the challenge of accurately measuring cell size during live-cell microscopy.

The simplest strategy to determine cell size from imaging data is to directly measure cellular area, and then calculate cell volume by assuming that cells are spherical, cylindrical, or ellipsoid. While this strategy is effective in many bacteria and yeast, it is inadequate for irregularly shaped animal cells. A straightforward extension of this strategy is to instead measure nuclear area, calculate nuclear volume assuming a spherical nucleus, and then rely on the known correlation between nuclear volume and cellular volume, *i.e.*, the fixed karyoplasmic ratio, to estimate cell size (Jorgensen *et al.*, 2007; Neumann and Nurse, 2007; Edens *et al.*, 2013; Levy and Heald, 2016). Two disadvantages of this approach are that the correlation between nuclear and total cellular size is not perfect – for example, human cells’ karyoplasmic ratios are lower in differentiated cells, and higher in high-grade tumors (Kumar *et al.*, 2015) – and that it is sensitive to small errors in nuclear area measurements. Nevertheless, it has been used successfully to demonstrate that cell cycle phase modulates cellular growth rate and to identify a role for p38 in maintenance of cell size homeostasis (Ginzberg *et al.*, 2018; Liu *et al.*, 2018).

Cell volumes can be reconstructed in three dimensions using confocal microscopy when the cell membrane is fluorescently labeled. However, this method is limited by phototoxicity in live-cell experiments, and relies on challenging 3D image segmentation (Bajcsy *et al.*, 2015). Another microscopy-based approach that measures cell dry mass is quantitative phase microscopy (QPM). This technique detects the phase shift as light waves pass through a specimen whose refractive index differs from that of the medium. The specimen’s optical path length can then be calculated by comparing the phase shift through the specimen to a blank reference image. Since the specimen’s optical path length is proportional to its dry mass, QPM provides a per-pixel optical mass. QPM can be difficult to use because it requires a specialized camera that only works with some objective lenses and its accuracy depends on acquiring a good reference image and precisely defining cell boundaries (Barer, 1952; Mir *et al.*, 2011; Zangle and Teitell, 2014).

Three novel techniques have recently expanded the repertoire for measuring cell size. Microchannel resonators contain a microfluidic channel suspended within a vibrating cantilever. When single suspended cells flow through the channel, they displace medium and the overall mass of the cantilever increases by the cell’s buoyant mass. This buoyant mass increase is then measured by the change in vibration frequency of the cantilever. By embedding microchannel resonators within a more complex microfluidic setup, it is possible to measure individual cells’ buoyant masses at multiple timepoints (Godin *et al.*, 2010; Son *et al.*, 2012, 2015; Cermak *et al.*, 2016). Furthermore, other techniques, such as fluorescence microscopy or single-cell RNA sequencing, can be added (Stevens *et al.*, 2016; Cetin *et al.*, 2017; Kimmerling *et al.*, 2018). The microchannel resonator approach is powerful and precise but requires specialized microfluidic instrumentation, is low-throughput, and is not easily applied to adherent cells (Park *et al.*, 2008, 2010). Another new technique is fluorescence exclusion microscopy in which cells are grown in medium containing fluorescent dextran. As long as the cells do not take up the fluorescent compound, the “background” fluorescence of a region can be measured. A cell displaces fluorescent media so that the amount of excluded corresponds to cell volume. This approach does require cells to grow in a specialized low-ceiling device (Bottier *et al.*, 2011; Zlotek-Zlotkiewicz *et al.*, 2015; Cadart *et al.*, 2018). Both microchannel resonators and fluorescence exclusion were used to show that cell density is nearly constant throughout the cell cycle, but that cells swell ∼20% during mitosis (Grover *et al.*, 2011; Bryan *et al.*, 2013; Son *et al.*, 2015; Zlotek-Zlotkiewicz *et al.*, 2015). A final option is to sidestep the difficulty of measuring irregular shapes by forcing animal cells to adopt more convenient morphologies. For example, cells can be grown in narrow channels requiring them to adopt a rod-shaped morphology, where length is directly proportional to volume (Varsano *et al.*, 2017). However, it is unknown to what extent such shape manipulation affects cell physiology.

Here, we present an alternative strategy, inspired by work in yeast (DiTalia *et al.*, 2007), that can be added to the toolkit of researchers seeking to measure animal cell size. Our approach relies on the fact that most proteins in a cell are produced proportionally to the total protein content of the cell (Figure 1B) (Newman *et al.*, 2006; Zhurinsky *et al.*, 2010; Neurohr *et al.*, 2019). In this report we show that a constitutively expressed nuclear fluorescent protein can easily be used to measure animal cell size.

## RESULTS

### Construction of a *prEF1α-mCherry-NLS* size reporter cell line

Good candidate promoters for a fluorescent total protein reporter should be highly, ubiquitously, and constitutively expressed. Promoters for genes involved in protein translation frequently meet these criteria. We selected the promoter of the translation elongation factor *EF1α* because it has also been commonly used in lentiviral infection systems (Chang *et al.*, 2011; Cheng *et al.*, 2011; Schwarz *et al.*, 2018). We used this promoter to drive expression of either of two nuclear localized fluorescent proteins: mCherry-NLS and E2-Crimson-NLS. We selected these fluorescent proteins due to their fast maturation, long lifespan, and compatibility with other fluorescent reporters of interest such as mAG-Geminin and an Rb-Clover fusion protein (Shaner *et al.*, 2004; Strack *et al.*, 2009). Localizing the fluorescent reporter to the nucleus facilitates image segmentation and measurement of total protein via wide field fluorescence imaging.

To construct a cell line with a total protein reporter, we inserted a *prEF1α-mCherry-NLS* expression cassette into immortalized human mammary epithelial cells (HMECs) by lentiviral infection and confirmed bright nuclear expression of the fluorescent protein (Figure 1C-D). Since we expected that expression variability due to gene copy number and location within the genome could be a major source of noise when comparing expression across cells, we sorted single cells by FACS and expanded clones.

Next, we set about assessing how well mCherry expression reflected cell size within a clone. Because our approach only works in cells, it cannot be validated by measuring non-cell objects of known sizes, such as beads. Therefore, since there is no single gold-standard method for size measurement (Table 1), we compared mCherry-NLS expression to several established methods.

### Constitutively expressed mCherry-NLS correlates with scatter, nuclear volume, and total protein

We incubated *EF1α-mCherry-NLS* HMECs with the protein dye CFSE and used flow cytometry to measure individual cells’ forward scatter (FSC), CFSE amount, and mCherry amount. We plotted each pair of measurements and performed a linear regression (Figure 2A-C). The intercepts for all three lines were close to the origin, indicating that all three measurements are approximately proportional. We found similar correlation coefficients (R^2^ between 0.4 and 0.6) between all three pairs of measurements, suggesting that no one measurement is substantially noisier than the others. We compared these cells to HMECs expressing mCherry-NLS from the *ACTB* promoter and determined that *EF1α-mCherry-NLS* was both brighter and more proportional to size (Figure S1A-B). To test whether our strategy also works in another cell type, we introduced *EF1α-mCherry-NLS* into K562 cells. We found that also in these cells, mCherry-NLS was proportional to FSC (Figure S1C).

**Figure 2.**
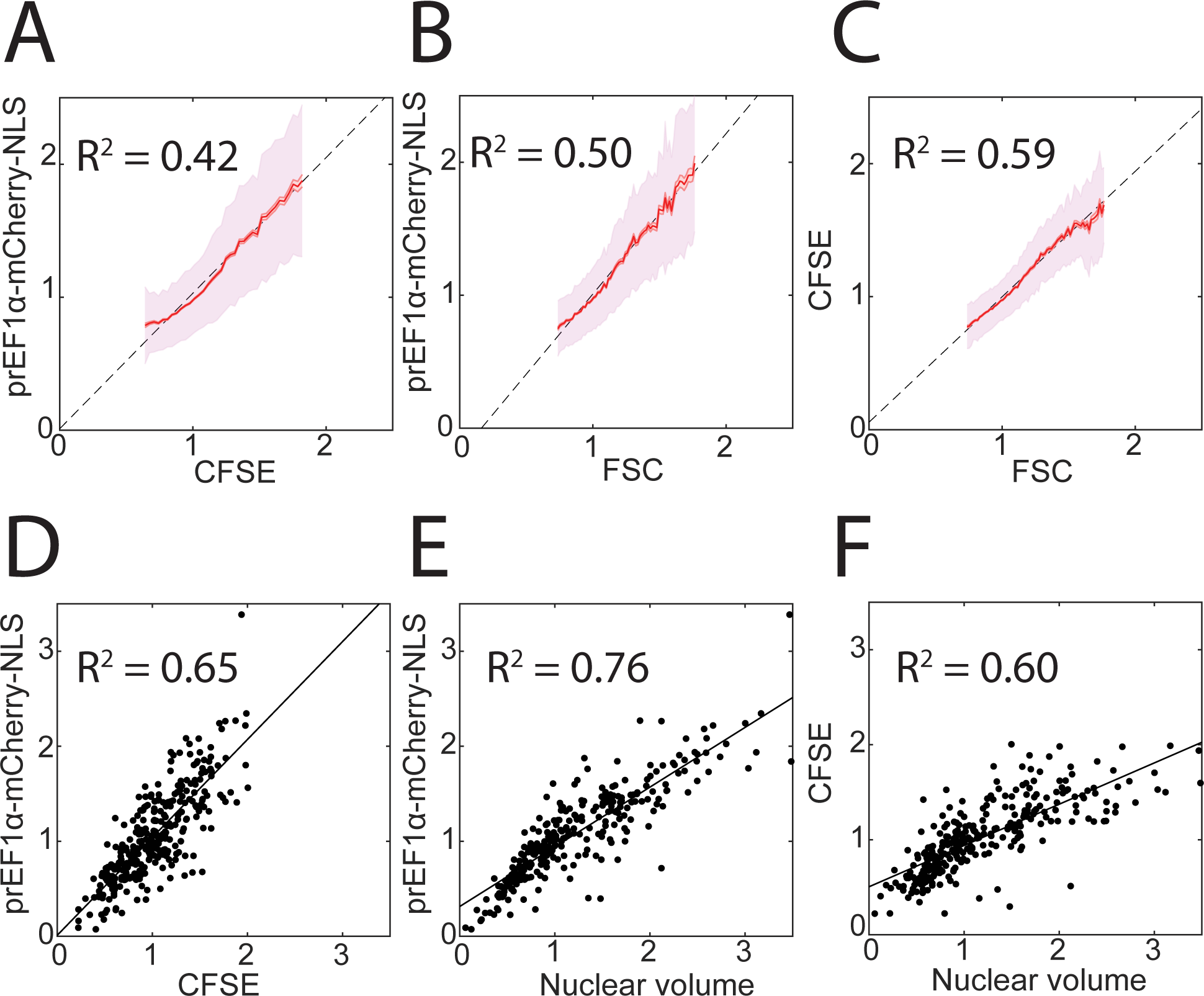
Comparison of constitutively expressed fluorescent protein with other cell size metrics. (A-C) Binned means and standard deviations for *prEF1α-mCherry-NLS*, CFSE, and FSC as measured by flow cytometry. CFSE is a total protein dye and FSC is the optical forward scatter (see methods). Each metric was normalized to its own median, and bins are shown for the middle 95% of the data. Least-squares regression lines and correlation coefficients are also shown. *N* > 30,000 cells. (D-F) Single cell measurements made using wide field fluorescence microscopy, least-squares linear regression, and correlation coefficients for the indicated metrics. *N* = 303 cells.

Next, we returned to *EF1α-mCherry-NLS* HMECs and examined them by fluorescence microscopy rather than flow cytometry. In this experiment, we measured nuclear volume (calculated as [nuclear area]^3/2^) and performed a pairwise comparison with mCherry-NLS and with CFSE total protein dye fluorescence signals (Figure 2D-F). Once again, we observed that all three intercepts were close to the origin and that the correlation coefficients were similar, suggesting that none of the three measurements is substantially inferior to the other two.

### Constitutively expressed mCherry-NLS size estimates are less sensitive than nuclear volume to variation in image segmentation

Since nuclear volume and mCherry-NLS total intensity are the two most experimentally straightforward approaches to measuring live cell size, we sought to compare them head-to-head. One pitfall of fluorescence microscopy is that it is necessary to distinguish foreground objects (cells or nuclei) from background by segmenting the image, often by applying a brightness threshold. We examined how varying our nuclear segmentation threshold (mCherry pixel intensity) affected our measurement of nuclear volume and mCherry-NLS total intensity. One representative nucleus segmented at different thresholds is shown in Figure 3A. Nuclear volume for this cell was highly sensitive to threshold choice, especially at less stringent thresholds, while mCherry-NLS total intensity was robust to threshold choice. To confirm this observation, we applied different thresholds to all the cells in our images and saw a similar pattern (Figure 3B).

**Figure 3.**
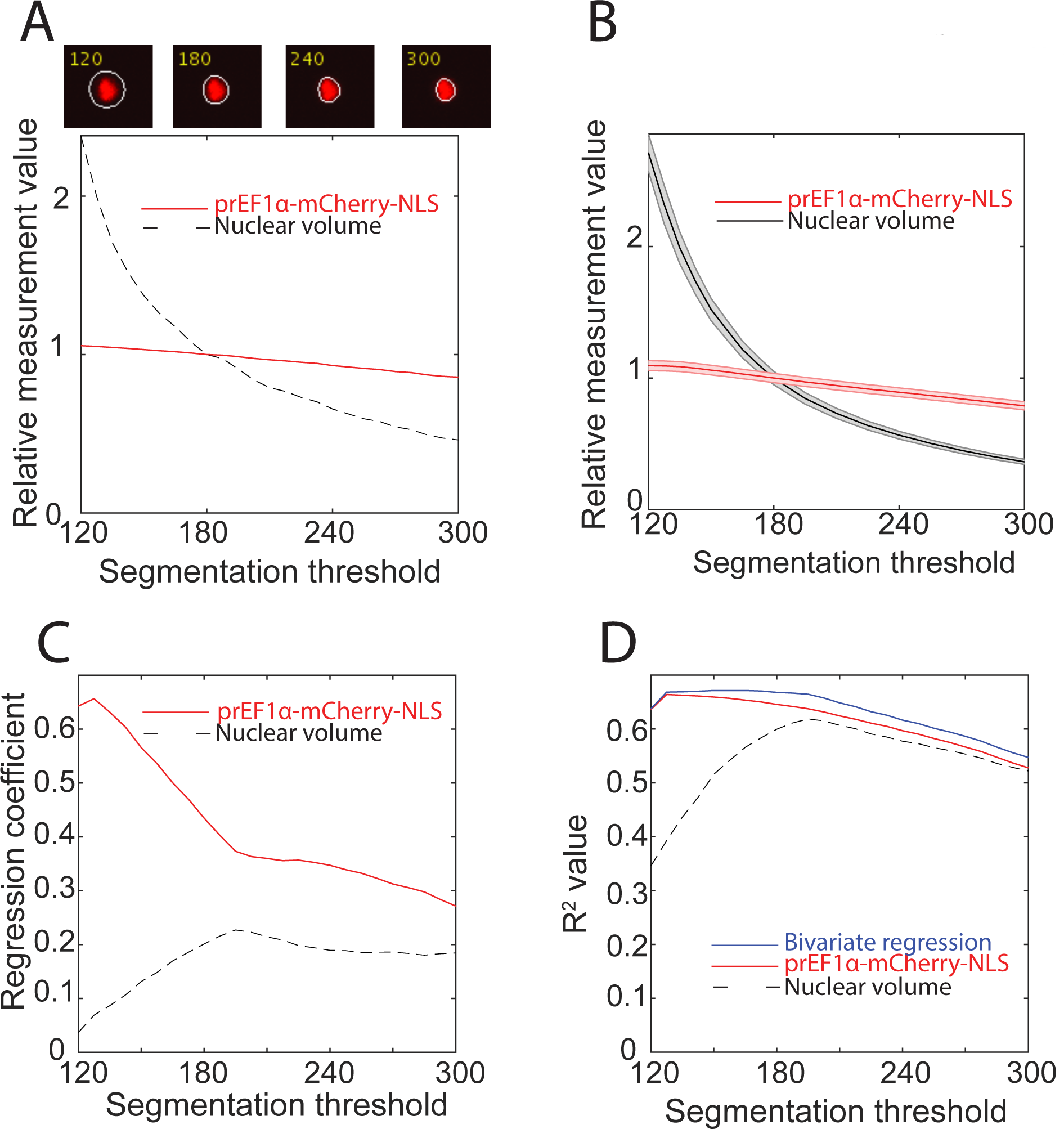
Sensitivity of nuclear volume and *EF1α-mCherry-NLS* to changes in segmentation threshold. (A) Upper panels: a single nucleus segmented at four different intensity thresholds. Lower panel: nuclear volume and total mCherry-NLS measurements for that cell at each threshold. Measurements were normalized to their value when the segmentation threshold is 180 (see methods). (B) Mean and associated standard error of nuclear volume and mCherry-NLS measurements for a cell population at different intensity thresholds. Measurements were normalized to the median value when the segmentation threshold is 180. *N* = 379 cells. (C-D) Analysis of correlation of nuclear volume and mCherry-NLS with CFSE total protein dye. Each measurement was normalized to its mean, and bivariate regression was performed using nuclear volume and mCherry-NLS as predictor variables, and using CFSE as the response variable. *N* = 303 cells. (C) Plot of the regression coefficient, *i.e*., slope, for each predictor variable as a function of segmentation threshold. (D) Plot of correlation coefficient as a function of segmentation threshold for the bivariate regression alongside the correlation coefficients for each single variable regression.

The reason for this phenomenon can be seen in the representative nucleus images. We can be confident that the region segmented at the most stringent threshold is truly part of the nucleus, while the peripheral halo that expands as the threshold decreases is less certain to be truly part of the nucleus. Adding or subtracting a single pixel from the uncertain periphery of the segmented area has an equivalently large effect on nuclear area or volume as adding or subtracting a pixel from the high-confidence central region. However, because the vast majority of the mCherry fluorescence is contained in the high-confidence central region, changes in the segmentation threshold have only a small effect on the total mCherry fluorescence measured. Thus, mCherry fluorescence is less sensitive than nuclear volume to changes in segmentation.

To further compare nuclear volume and mCherry fluorescence, we stained cells with the total protein dye CFSE and again segmented the nuclei at varying thresholds. At each threshold, we performed a bivariate regression with CFSE as the response variable and nuclear volume and mCherry as the predictor variables. Both nuclear volume and mCherry were normalized to their mean values. The regression coefficient for the mCherry protein reporter was higher than for nuclear volume, indicating that CFSE intensity was predominately predicted by our protein reporter. This effect is enhanced at less stringent segmentation thresholds (Figure 3C). Moreover, adding nuclear volume to the regression only marginally improves the correlation with CFSE compared with mCherry alone (Figure 3D). Taken together, these data support preferential use of our *EF1α-mCherry-NLS* construct over nuclear volume to report total cellular protein content.

### Constitutively expressed mCherry-NLS measurements are less noisy than nuclear volume measurements in live cells tracked over time

Having established that constitutively expressed mCherry-NLS reflects size in populations of cells at a single moment in time, we turned our attention to live cell microscopy to examine size dynamics in single cells. We imaged *EF1α-mCherry-NLS* HMECs for 48-72 hours, segmented the nuclei, and tracked them over time. We examined growth of single cells by looking at single cell traces of nuclear volume or mCherry intensity over time (Movie 1; Figure 4A-B). We then compared variability of nuclear volume versus mCherry total intensity within single cell traces. Since nuclear volume and total protein content are only expected to change slowly over time, variation in the signal over shorter time scales is presumably due to measurement and/or segmentation error. To estimate the magnitude of these sources of error, we approximated each trace with overlapping piecewise linear fits and summed the squared residuals from each timepoint (Figure 4C-D). We found significantly more timepoint-to-timepoint variability in nuclear volume than in mCherry total intensity (Figure 4E). As in Figure 3, this result is because small changes in image segmentation have a large effect on nuclear volume but not total mCherry measurements.

**Figure 4.**
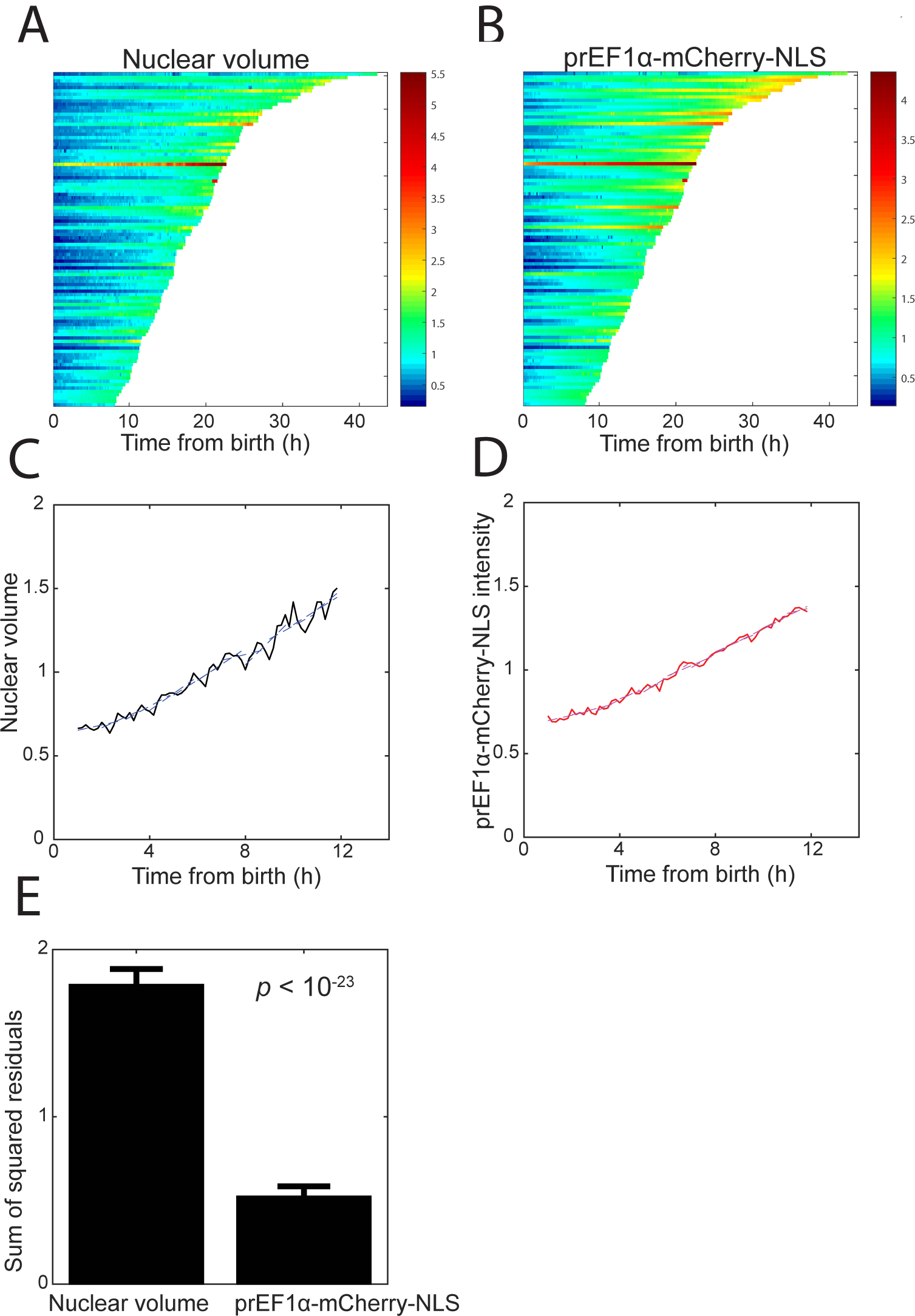
Nuclear volume and mCherry-NLS size measurements for live single cells tracked through time. (A-B) Single cell traces over time. Measurements were normalized to the mean of all the data. (C-D) Representative single cell traces along with a piecewise fit to quantify timepoint-to-timepoint variability. Each trace was normalized to its own mean. (E) Plot of an error metric: the mean and standard error of the sum of squared residuals from the piecewise linear fits to each cell as in (C-D).

### Constitutively expressed mCherry-NLS correlates with dry mass measurements

Next, we sought to compare our mCherry-NLS reporter with quantitative phase microscopy that measures cell dry mass. In live cells tracked through time, we measured dry mass, mCherry total intensity, and nuclear volume. We found that mCherry-NLS and dry mass were less noisy than nuclear volume (Figure 5A-B). Note that because accurate automated segmentation (as in Figure 4) was not possible in these quantitative phase images, all three measurements for this experiment are from manual segmentation. When we compared measurements across multiple cells, dry mass correlated better with mCherry than with nuclear volume. However, unlike with CFSE, these curves do not intercept close to the origin, highlighting the challenge of precisely taring quantitative phase data (Figure 5C-D). Based on these experiments and those described in the previous section, we concluded that mCherry intensity continues to reflect cell size during live cell imaging.

**Figure 5.**
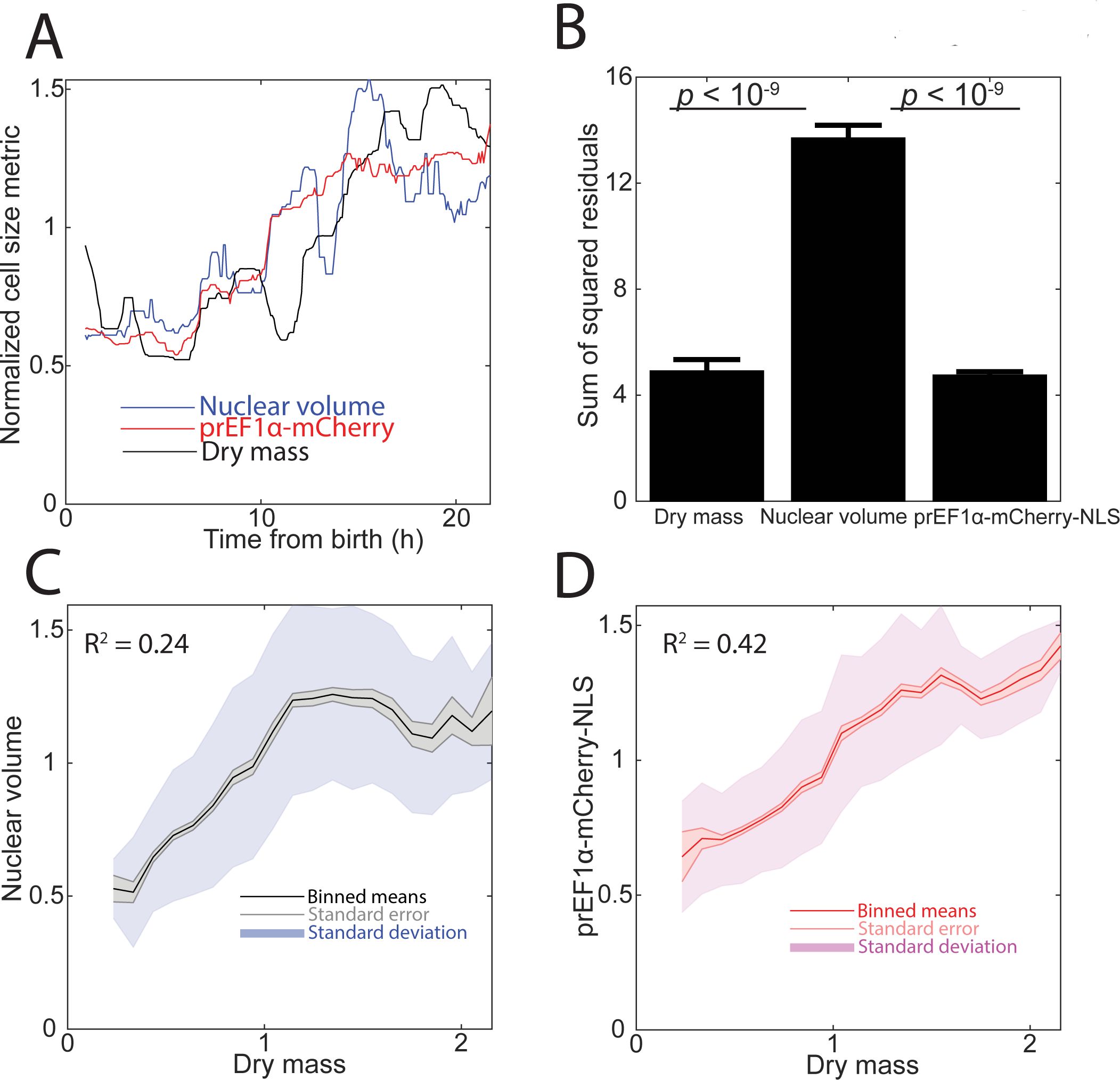
Comparison of nuclear volume, mCherry-NLS, and dry mass measurements. Dry mass was measured by quantitative phase microscopy (see methods). (A) Representative trace of a single cell tracked through time. Each measurement type was normalized to its median. The decrease in dry mass around 11 hours after birth is an artifact of reference background subtraction (see methods). (B) Plot of an error metric: the mean and standard error of the sum of squared residuals from a piecewise linear fit, as in Figure 4C-D. *N* = 8 cells. (C-D) Correlation of nuclear volume or mCherry-NLS with cell dry mass. Binned means, standard error of the mean, and standard deviation of the data are shown. *N* > 2000 measurements.

### Constitutively expressed E2-Crimson-NLS enables measurement of G1 size control

Finally, we applied our protein content measurements to describe cell growth and the concentration dynamics of key cell cycle regulators through the cell cycle, which we anticipate will be a frequent application of our method. To show how our size reporter can be used to measure concentration dynamics of a key cell cycle regulator through the cell cycle, we integrated our size reporter into cells expressing the G1/S inhibitor protein RB (retinoblastoma protein) tagged with Clover green fluorescent protein from its endogenous locus. *RB* was tagged using CRISPR/Cas9 as described in (Zatulovskiy *et al.*, 2018). We infected *RB-Clover* cells with lentivirus particles containing a Geminin-mCherry reporter construct to define the G1 to S phase transition by the accumulation of red fluorescence (Sakaue-Sawano *et al.*, 2008). Due to using a red fluorophore as a cell cycle phase reporter, we switched to far-red *prEF1α-E2-Crimson-NLS* to measure size (Figure 6A). This color-swapped reporter also correlates well with other size metrics (Figure S2). We were able to track cells and make fluorescence measurements in 3 colors without phototoxicity (Figure 6B; Figure S3). Rb protein concentration can then be measured as the ratio of Clover to E2-Crimson-NLS total fluorescence and plotted as a function of cell cycle phase (Figure 6C). We observed that Rb concentration decreases during G1 and increases again starting in S phase, consistent with (Zatulovskiy *et al.*, 2018).

**Figure 6.**
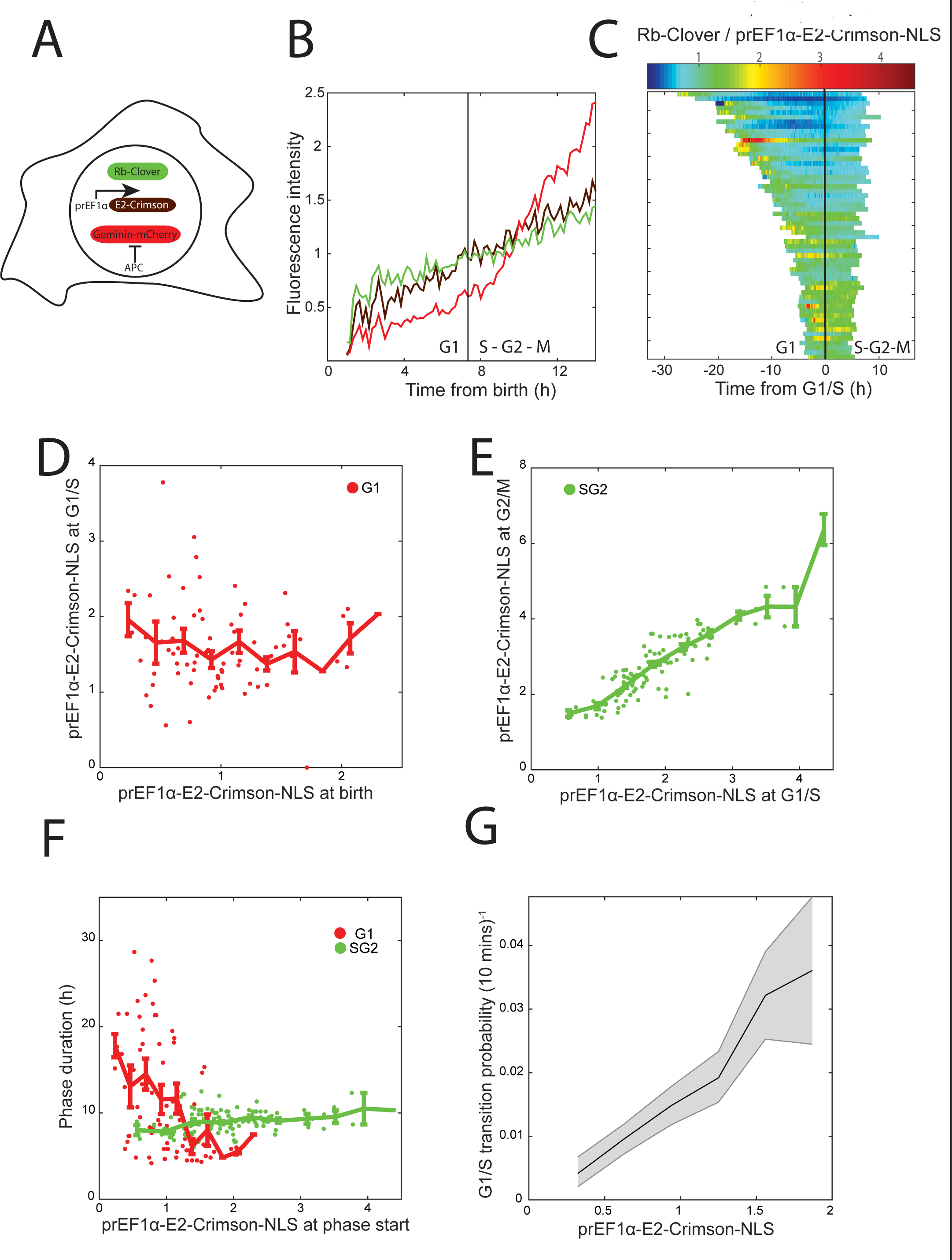
Using a constitutively expressed size reporter to analyze the relationship between cell size and cell cycle progression. (A) Schematic of the three fluorescent reporters introduced into each cell. Rb-Clover fluorescent fusion protein was expressed from one of the two endogenous *RB* alleles. A Geminin-mCherry reporter was used to identify the G1/S transition (see methods). (B) Representative trace of the three fluorescent reporters from cell birth to mitosis. Each measurement type was normalized to its mean. (C) Plot of Rb protein concentration (calculated as Rb-Clover amount divided by E2-Crimson-NLS amount). Cells were aligned at the time of their G1/S transitions. (D) Plot of the amount of E2-Crimson-NLS at birth versus the amount at G1/S. Data were normalized to the mean amount of E2-Crimson-NLS at birth and are shown with binned means ± standard error of the mean. (E) Plot of the amount of E2-Crimson-NLS at G1/S versus the amount at G2/M. Data were normalized to the mean amount of E2-Crimson-NLS at birth and are shown with binned means ± standard error of the mean. (F) Plot of the duration of G1 or S/G2 phases versus E2-Crimson-NLS amount at birth or at G1/S respectively. *N* = 77 and 122 cells for G1 and S/G2 durations respectively. (G) Calculation of the rate at which cells stochastically pass the G1/S transition as a function of E2-Crimson-NLS amount (see methods). The fraction of measurements in each E2-Crimson-NLS bin that correspond to a G1/S transition at that frame is shown, along with 5^th^ and 95^th^ percentile confidence intervals from bootstrap resampling of each bin 1000 times. Bins are shown for the middle 90% of size measurements and were normalized to the overall mean. *N* = 290 transition events from >18,000 measurements.

It has previously been observed in several mammalian cell lines that cells adjust the amount of time they spend in G1 phase to compensate for differences in birth size, while the amount of time spent in S/G2 phases is unaffected (Cadart *et al.*, 2018). To investigate size control at G1/S in HMEC cells, we measured E2-Crimson total intensity ∼1 hour after birth (to allow time for nuclear reimport of E2-Crimson), at the G1/S transition as determined using Geminin-mCherry, and ∼1.5 hours before cytokinesis. We observed that smaller-born cells grow more during G1 than larger-born cells so that all cells are a similar size at the G1/S transition (Figure 6D). Conversely, during S/G2, cells add a similar amount of mass regardless of their size at G1/S so that cell size at mitotic entry was highly correlated with their size at G1/S (Figure 6E). We further observed that G1 length is negatively correlated with size at birth, while S/G2 length is remarkably constant (Figure 6F). As predicted by this model, we observed that larger cells enter S phase at a higher rate than smaller cells (Figure 6G). These results highlight the importance of G1 for maintaining cell size homeostasis, consistent with other reports (Killander and Zetterberg, 1965; Varsano *et al.*, 2017; Ginzberg *et al.*, 2018; Liu *et al.*, 2018; Zatulovskiy *et al.*, 2018).

## DISCUSSION

Here, we show that a constitutively expressed fluorescent protein provides a convenient metric of cell size. We validated this method by showing that integrated fluorescence intensity correlates well with more established size measurements including forward scatter from flow cytometry, CFSE total protein dye, nuclear volume, and quantitative phase microscopy measurements of dry mass. Importantly, we found that fluorescent reporter total intensity, nuclear volume, CFSE, and forward scatter were not only linearly related, but proportionally related so that the intercepts are close to the origin. While a linear relationship would mean that any metric could be used to rank cells in order of size, a proportional relationship implies that any can be used also to identify fold-differences in sizes between cells. We further showed that fluorescence intensity is less sensitive to segmentation differences and less noisy than nuclear volume. Finally, we measured cell size over time to recapitulate several prior observations: that Rb protein concentration falls during G1, that time spent in G1 is negatively correlated with birth size, that time spent in S/G2 is uncorrelated with cell size, and that larger cells have a higher rate of progression through the G1/S transition.

Our method is limited by several factors. First, it requires generating a clonal cell population expressing the fluorescent reporter of choice. Second, it occupies a fluorescence channel and therefore may not be compatible with experiments in which more than two or three other color channels are being used. Third, it relies on the scaling of individual proteins with cell size. While this principle has been consistently demonstrated in the normal cell size range, it may not work for abnormally large cells (Neurohr *et al.*, 2019). Finally, our method provides only a relative, not an absolute, measurement of cell size. However, by combining our method with an orthogonal approach to measure the average absolute size of the population, it would be possible to calculate individual absolute cell sizes. Despite these limitations, we expect that our straightforward strategy for measuring cell size will enable other investigators to routinely ascertain how size and growth affect other areas of cell biology. Importantly, the combination of our size reporter with another fluorescently labeled protein allows the measurement of that protein’s concentration using a common wide field microscope. This could be especially impactful for studies determining how dynamic protein concentrations control key cellular events.

## Materials and Methods

### Cell culture conditions and cell lines

Cells were cultured at 37°C with humidified 5% CO_2_. Non-transformed hTERT1-immortalized human mammary epithelial cells (HMECs) were obtained from the laboratory of Stephen Elledge at Harvard Medical School (Solimini *et al.*, 2012) and cultured in MEGM Mammary Epithelial Growth Medium (Lonza CC-3150). To reduce background fluorescence in imaging experiments we used the same medium without phenol red (Lonza CC-3153 basal medium supplemented with growth factors and other components from the CC-4136 kit). Medium was refreshed every 24 hours during imaging experiments to mitigate phototoxicity and nutrient depletion. Endogenously tagged *RB-Clover* cells were generated as described in (Zatulovskiy *et al.*, 2018). HEK 293T cells (for producing lentivirus) were cultured in Dulbecco’s modified Eagle’s medium (DMEM) containing L-glutamine, 4.5 g/L glucose, and sodium pyruvate (Corning), supplemented with 10% fetal bovine serum (Corning) and 1% penicillin/streptomycin. K562 cells were a gift from Ravi Majeti’s lab at Stanford University and were generally cultured in RPMI 1640 medium (HyClone) supplemented with 10% fetal bovine serum and 1% penicillin/streptomycin. For 3 weeks after single-cell sorting, K562 clones were expanded in IMDM medium (Gibco) supplemented with 20% fetal bovine serum and 1% penicillin/streptomycin.

### Fluorescent reporter cell lines

Fluorescent reporters (mCherry-3xNLS, E2-Crimson-NLS, mAG-hGeminin, and mCherry-hGeminin) were cloned into the CSII-EF-MCS lentiviral vector backbone as in (Schwarz *et al.*, 2018). The CSII vector, the lentiviral packaging vector dr8.74, and the lentiviral envelope vector VSVg were transfected into HEK 293T cells with TurboFect (Life Technologies). Transfected HEK 293T cells were incubated for two days before the virus-laden medium was transferred to HMECs or K562 cells. At least 3 days after infection, single cells positive for the fluorescent protein of interest were sorted by FACS and clonally expanded.

### Cell imaging

Cells were seeded the day before imaging on 35mm glass-bottom dishes (MatTek) at low density. The cells were kept overnight in standard culture conditions before being transferred to a Zeiss Observer Z1 microscope equipped with an incubation chamber to maintain the cells at 37°C with humidified 5% CO_2_. For live cell experiments, fluorescence and brightfield or phase images were collected every 10 minutes for 48-72 hours using a Zyla 5.5 sCMOS camera, a 10X Plan-APOCHROMAT 0.45 NA or 10X Plan-NEOFLUAR 0.30 NA objective, a Colibri.2 light source, and an automated stage, all controlled by μManager software (Edelstein *et al.*, 2014). This imaging setup has a thick enough focal plane to collect emitted light from the entire nucleus without z-stacks. For total protein dye experiments, cells were incubated with CFSE (Thermo Fisher C34554) at 1:2500 dilution for 15 minutes at 37°C. We did not observe significant photobleaching during our experiments (Figure S4).

### Flow cytometry and cell sorting

HMECs were grown on dishes to ∼70% confluence and harvested by trypsinization and centrifugation, then resuspended in PBS. To generate clonal cell lines, cells were sorted using a BD FACSAria machine. For total protein dye experiments, cells were incubated with CFSE (Thermo Fisher C34554) at 1:4000 dilution for 15 minutes at 37°C. K562 cells were harvested by centrifugation and resuspended in PBS. For all analyses, flow cytometry was performed using a BD LSRII.UV instrument, then single cells were gated using FlowJo. Forward scatter area (FSC-A), CFSE area, and mCherry or E2-Crimson area were each normalized to the population median. For clarity, bins are shown for the middle 95% of the data due to wide variance and nonlinearities in measurements for cells of extreme sizes. Plots were made with the MATLAB function “shadedErrorBar”.

### Fluorescence image analysis

Images were quantified using custom MATLAB scripts (Mathworks). In MATLAB, cell nuclei were segmented using the mCherry or E2-Crimson channels by Gaussian filtering, thresholding with manually chosen parameters, and opening and closing of the segmented regions using the MATLAB functions imopen and imclose. Nuclear volume was calculated as the segmented nuclear area (in pixels) to the 3/2 power. We also performed our analyses using nuclear area rather than nuclear volume with qualitatively similar results. For CFSE experiments, complete cells were similarly segmented using the CFSE channel, and additional thresholds were applied to omit cell debris from subsequent analysis. For all imaging experiments, total pixel intensity in the segmented region was calculated after applying a location-dependent intensity adjustment to account for vignetting due to our large camera field of view, which caused the center of an image to be illuminated more brightly than the periphery. To make the adjustment we used an image of dissolved fluorescein in solution to measure effective illumination as a function of position in the field of view. Then, each pixel in each image of cells was divided by the difference between the effective illumination intensity and the darkfield intensity at that pixel location (Figure S5), as in (Bottier *et al.*, 2011). Finally, each segmented object’s local background was subtracted to account for fluorescence of the cell medium.

### Segmentation threshold analysis

A range of possible thresholds that yielded plausible nuclear segmentations was manually chosen, and nuclei were automatically segmented at each possible threshold. A threshold of 180 was judged to be the closest to manual segmentation and was therefore used as the reference point. Nuclear volume and mCherry total intensity were calculated at each possible threshold and normalized to their respective values for the cell of interest at the reference threshold (for Figure 3A) or to the mean value for all cells at the reference threshold (for Figure 3B). For regression analysis, each metric was normalized to its own mean. At each possible threshold, uni- and bivariate regression was performed using either nuclear volume, mCherry total intensity, or both as predictor variables and CFSE total intensity as the response variable. The regression coefficients in Figure 3C represent the change in normalized CFSE total intensity per unit change in normalized nuclear volume or mCherry intensity. The squared correlation coefficients in Figure 3D represent the goodness of fit for the bivariate regression compared with each univariate regression.

### Live cell imaging analysis

Nuclei were manually tracked over time with the aid of a custom MATLAB graphical user interface. During tracking, in cases where automated segmentation failed to separate neighboring nuclei, the fused object was split by applying a watershed algorithm while decreasing the size of the morphological structuring element until the desired number of distinct nuclei were segmented. During subsequent analysis, additional noise, *e.g.*, due to segmentation errors or nearby cell debris, was corrected as follows. At timepoints where the cell was not segmented at all, the average of the measurements at the timepoints before and after the missing timepoint was used. Then, the measurement at each timepoint was compared to a moving median with window size equal to nine frames. If the difference between the actual measurement and the moving median was more than half the magnitude of the moving median, the moving median was used instead. Finally, measurements from the first hour after birth were omitted on account of low-confidence segmentation while the cells reimported fluorescent protein into their postmitotic nuclei, and measurements from the last two hours before cytokinesis were omitted on account of nuclear envelope breakdown. For analysis of timepoint-to-timepoint measurement noise, each trace was normalized to its own mean and then decomposed into an overlapping series of two-hour segments. A line was fit to each two-hour segment. Then, the residuals were calculated for each segment, squared, and summed across all segments. The resulting sum of squared residuals was calculated for each nuclear volume or mCherry intensity trace, and this noise metric was compared for the two measurement types using a two-sided t-test.

### Quantitative phase microscopy

Quantitative phase microscopy was performed with assistance from Phasics (Saint-Aubin, France). Images were acquired using a wavefront SID4Bio camera using Quadriwave Lateral Shearing Interferometry, then processed using the manufacturer’s SID4Bio software according to the procedures in (Popescu *et al.*, 2008; Aknoun *et al.*, 2015). Briefly, a quadratic fit was applied to an empty reference frame and this fit was subtracted from the image being analyzed. These images were imported into ImageJ and manually segmented and tracked. Then, each object’s local background was subtracted. For these experiments, a 20X Plan-APOCHROMAT 0.8 NA objective lens was used and images were collected every 5 minutes. Since manual segmentation resulted in significant timepoint-to-timepoint variability, Figure 5A shows moving medians with a window size of 17 frames (∼1.5 hour). As in Figure 4C-E, for Figure 5B we performed a piecewise linear fit to the raw data, but used overlapping one-hour rather than two-hour segments to address noise in the data as described above. The sum of squared residuals for each measurement type were compared by ANOVA and a multi-comparison t-test. The correlations shown in Figures 5C-D were calculated by treating each single-cell, single-timepoint measurement as an independent data point.

### Cell cycle analysis

The Rb-Clover, E2-Crimson-NLS, and Geminin-mCherry traces shown in Figure 6B were normalized to their means for that trace so that they could be plotted on the same axes. The G1/S transition was automatically detected by an algorithm that looked for surpassing a threshold level of Geminin-mCherry, a kink in the Geminin-mCherry signal, and a local maximum of the second derivative of the signal. Because E2-Crimson fluorescence was also excited by the 555 nm LED used for mCherry fluorescence, in order to facilitate distinguishing the accumulation of Geminin-mCherry beginning at G1/S from constitutive accumulation of the E2-Crimson size reporter, the Geminin-mCherry trace was divided by the E2-Crimson trace so that the mCherry signal is flat in G1, where there is expected to be no stable Geminin protein. The raw Geminin-mCherry values and the G1/S transition point identified by this method are shown in Figure 6B. Birth size was defined as the median size between 60 and 120 minutes after birth (to allow time for cells to reimport fluorescent protein into their postmitotic nuclei), while G1/S size was defined as the median size in the 50 minutes surrounding G1/S, and G2/M size was defined as the median size in the 60 to 120 minutes preceding cytokinesis (to avoid complications from nuclear envelope breakdown). Rb-Clover concentrations per unit nuclear volume and per unit E2-Crimson were calculated as Rb-Clover total intensity divided by the median value of the appropriate size metric within a 5-frame window. For plotting the single cell traces, Rb-Clover concentrations were normalized to the mean concentration for all measurements across all cells. For the G1/S rate experiment, automated segmentation and tracking with Aivia software (DRVision Technologies) were used to supplement manually tracked cells. Total E2-Crimson intensity was measured for all cells at all timepoints up through the G1/S transition, and each measurement was annotated with 0 or 1 for whether G1/S occurred there. Measurements were binned by E2-Crimson intensity and the fraction of measurements within that bin corresponding to a G1/S transition was calculated, along with 5th and 95th percentile bootstrapped confidence intervals.

## Author Contributions

DFB, EZ, and JMS designed the study. DFB and EZ performed the flow cytometry experiments. DFB performed the imaging experiments. DFB wrote the analysis software with assistance from SX. DFB and JMS wrote the manuscript.

## Acknowledgements

We thank members of the Skotheim lab for discussion and comments on the manuscript. We thank Flor Medina, Valentin Genuer, and Sherazade Aknoun from Phasics for the quantitative phase microscopy measurements. We thank Madhav Mani from Northwestern University for helpful suggestions on image analysis. We thank Anna Rajaratnam for assistance with cell tracking. We thank the Elledge laboratory at Harvard Medical School for the HMEC cell line and the Majeti laboratory at Stanford for the K562 cell line. We thank Nicholas Buchler, Stefano Di Talia, and Joseph Lipsick for helpful comments on the manuscript. This work was supported by the NIH through R01 GM115479 and T32 GM007365 (DFB). Cell sorting and flow cytometry analysis for this project were done on instruments in the Stanford Shared FACS Facility: flow cytometry data were collected on the LSRII.UV instrument, and sorting was performed on the Falstaff instrument, both of which were obtained using NIH S10 Shared Instrument Grants (S10RR027431 and S10RR027431, respectively).

---

**Figure S1.**
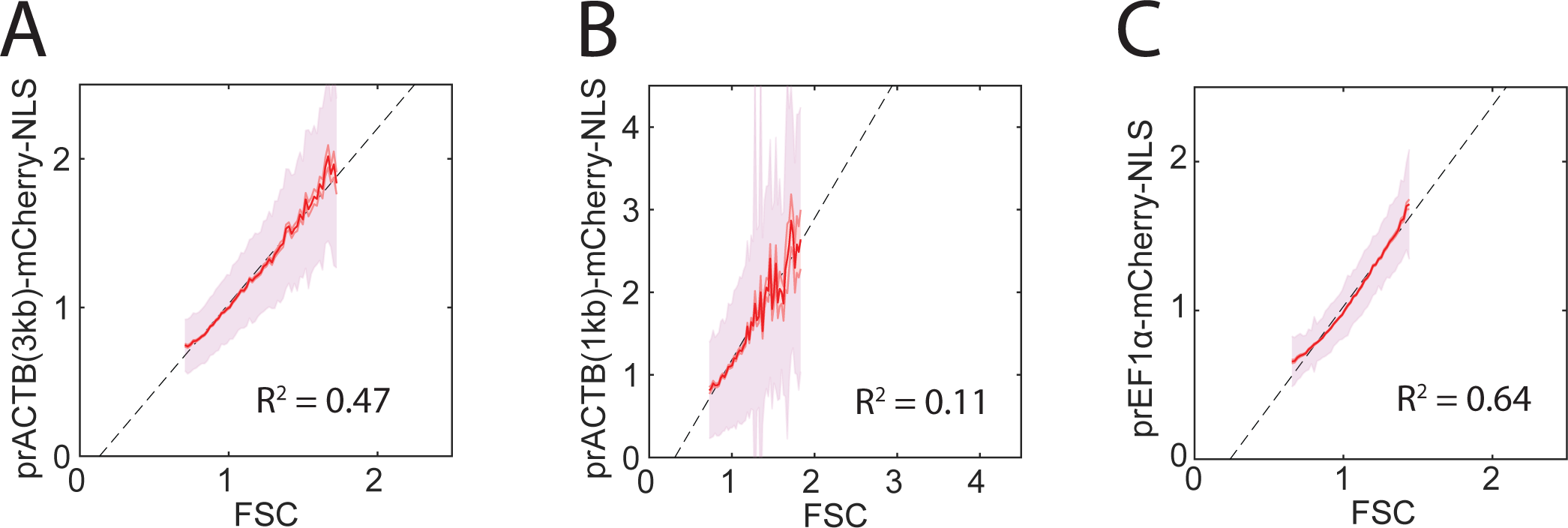
Comparison of mCherry-NLS with scatter when expressed from alternative promoters or in other cell lines. Binned means and standard deviations are shown for the middle 95% of the data after normalizing each metric to its own mean. Least-squared regression lines and correlation coefficients are also shown. *N* > 15,000 cells. (A) 3kb of the *ACTB* promoter was used to express mCherry-NLS in HMECs. (B) 1kb of the *ACTB* promoter was used to express mCherry-NLS in HMECs. (C) The *EF1α* promoter was used to express mCherry-NLS in K562 cells.

**Figure S2.**
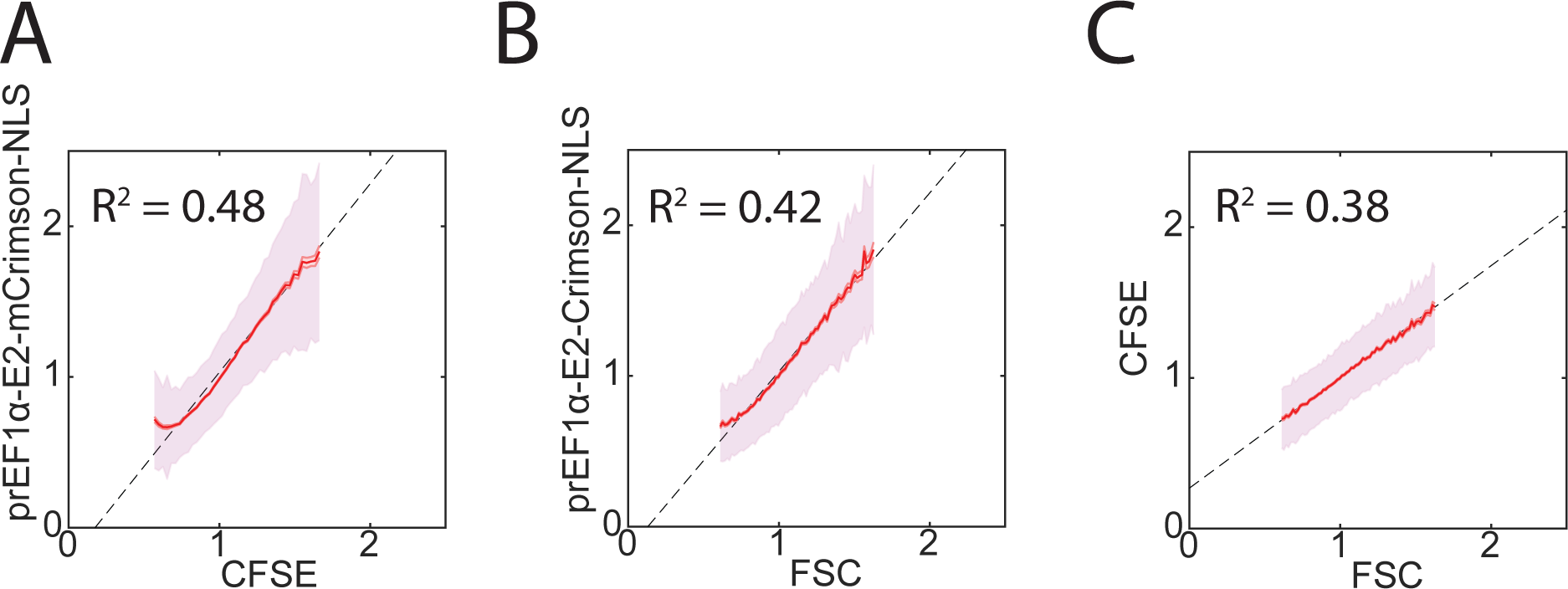
Comparison of constitutively expressed E2-Crimson with other cell size metrics. (A-C) Binned means and standard deviations for *prEF1α-E2-Crimson-NLS*, CFSE, and FSC as measured by flow cytometry. Each metric was normalized to its own median, and bins are shown for the middle 95% of the data. Least-squares regression lines and correlation coefficients are also shown. *N* > 50,000 cells.

**Figure S3.**
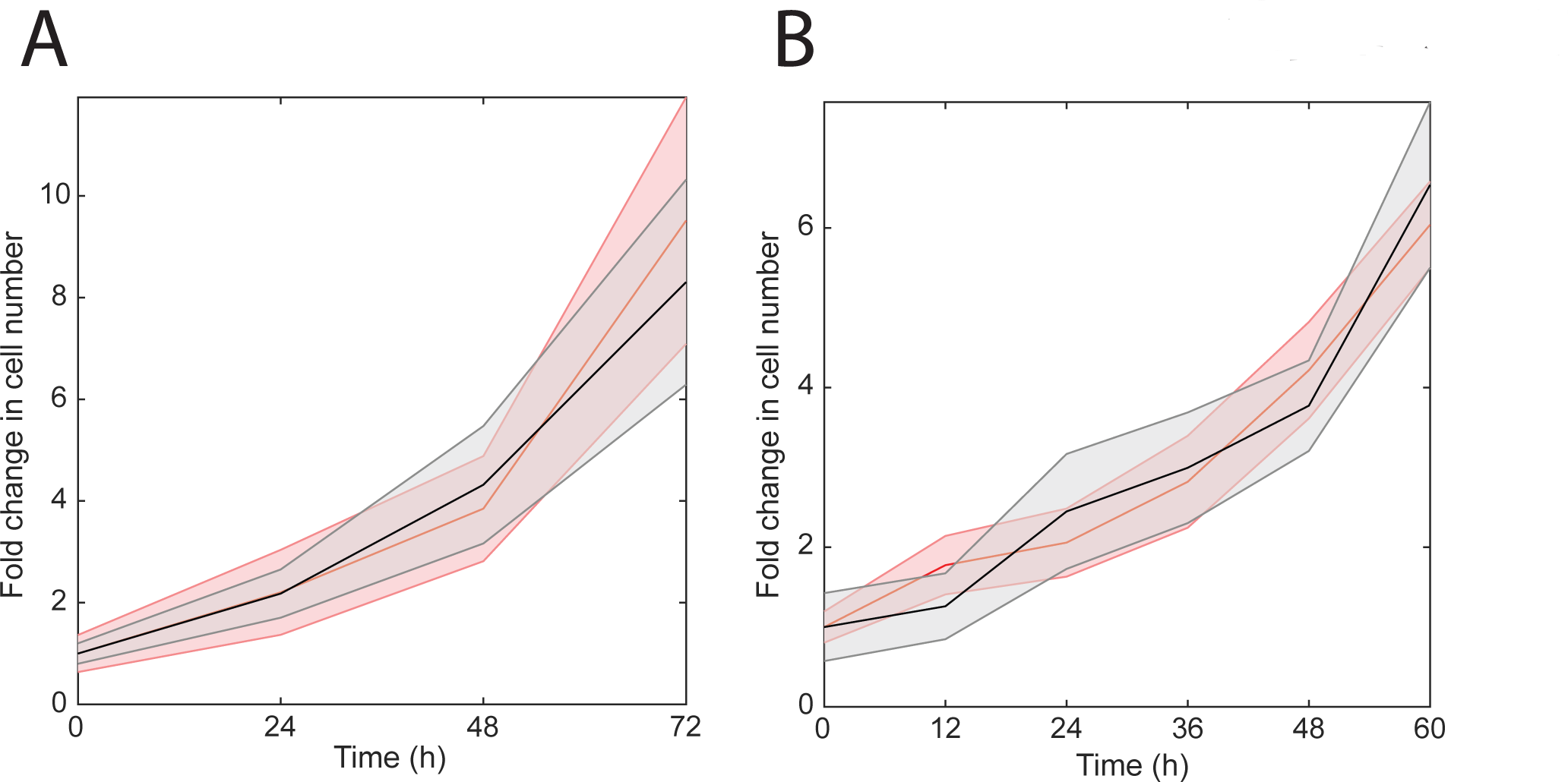
Growth curve for *prEF1α-mCherry-NLS* (A) or *prEF1α-E2-Crimson-NLS* (B) HMECs cultured in the microscope incubator under fluorescence imaging conditions (red) or transmitted light only (black). Cells were sparsely seeded and the numbers of cells in eight randomly selected fields of view were counted every 24 hours (A) or 12 hours (B), then normalized to the average number of cells per field at experiment start. Mean fold change is shown +/- standard error of the fold increase. There was no statistically significant difference between the two conditions in the fold increases at any timepoint (two sided t-test, *p* > 0.3).

**Figure S4.**
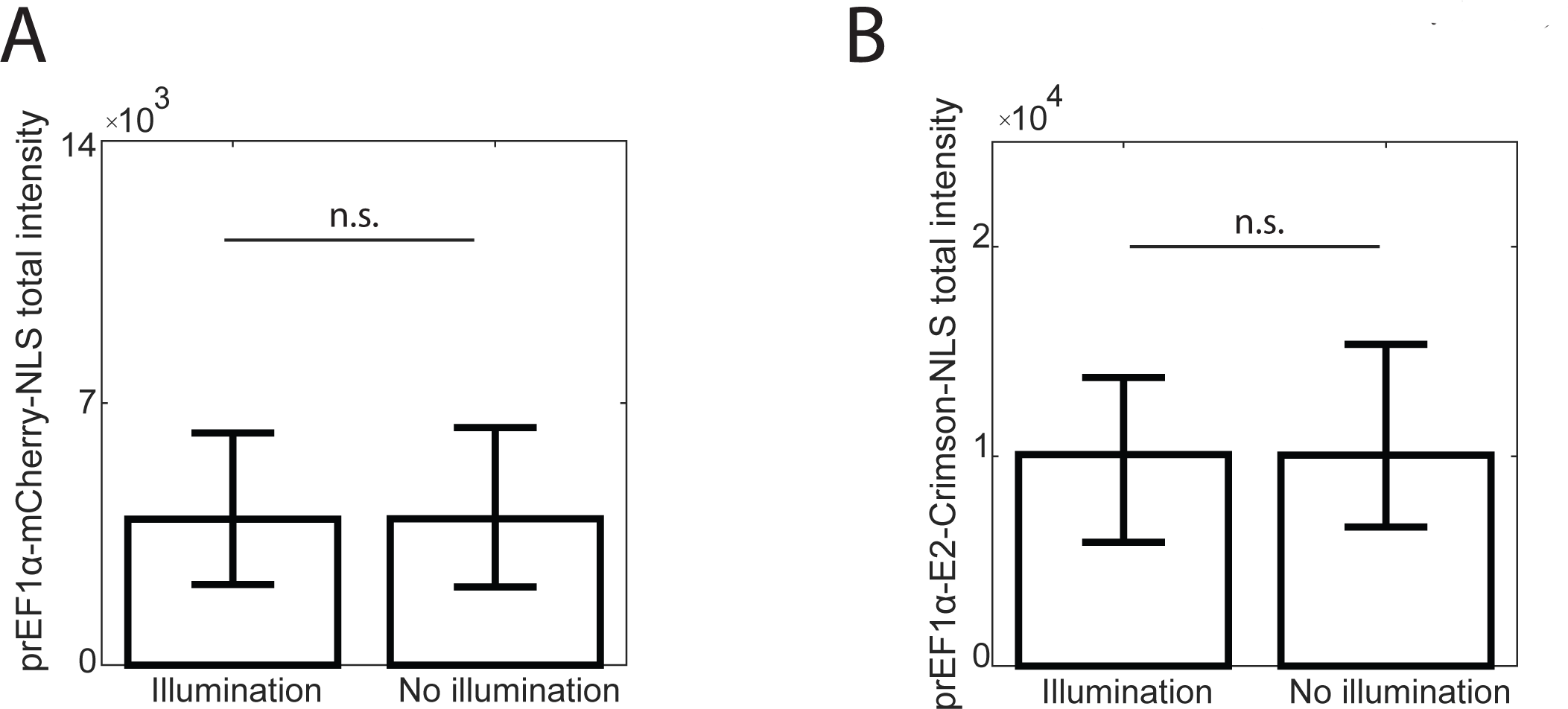
Effect of photobleaching on constitutively expressed fluorescent size reporter. (A) Cells expressing *prEF1α-mCherry-NLS* were imaged for 20 hours under fluorescence imaging conditions or without fluorescence illumination. After 20 hours, one additional fluorescence image was taken for each condition. The mCherry-NLS total intensity per cell was quantified automatically and is plotted as the median of all cells with the first and third quartiles. (B) Cells expressing *prEF1α-E2-Crimson-NLS* were imaged for 72 hours under fluorescence imaging conditions or without fluorescence illumination. After 72 hours, one additional fluorescence image was taken for each condition. The Crimson-NLS total intensity per cell was quantified manually and is plotted as the median of all cells with the first and third quartiles. For both (A-B), there was no statistically significant difference between the two conditions (two-sided t-test, *p* > 0.1).

**Figure S5.**
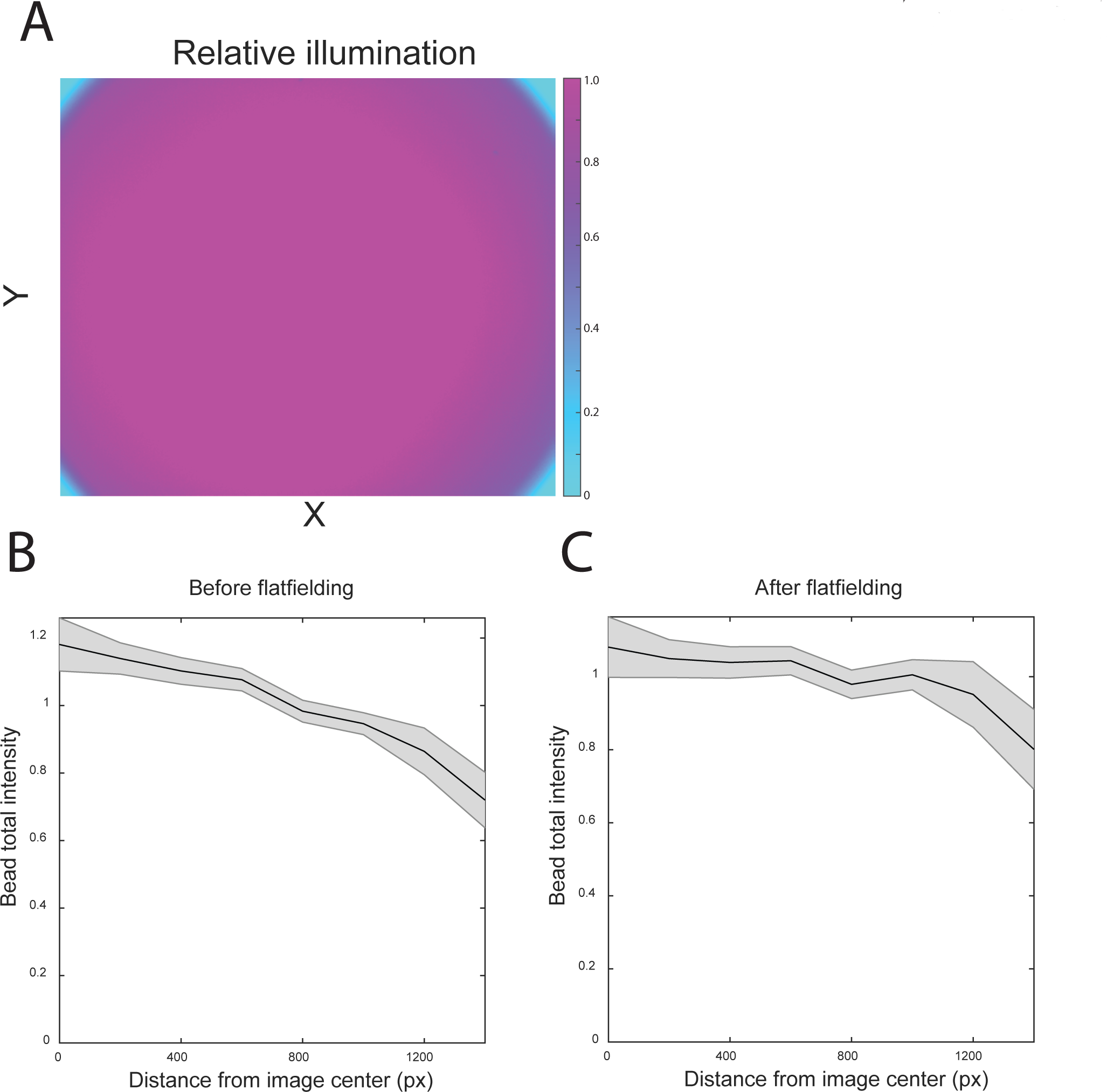
Correction of uneven illumination. (A) Pseudocolor image of fluorescein dissolved in PBS at 5μg/mL. In addition to the complete lack of illumination at the corners, decreased illumination of the periphery of the image relative to the center can be observed. (B) TetraSpeck beads were imaged and their total fluorescence intensity was plotted against their distance from the image center. (C) The same data as in (B), but after the flatfielding correction was applied as in (Bottier *et al.*, 2011).

